# Mex67 paralogs mediate division of labor in trypanosome RNA processing and export

**DOI:** 10.1101/2022.06.27.497849

**Authors:** Samson O. Obado, Milana Stein, Eva Hegedűsová, Wenzhu Zhang, Sebastian Hutchinson, Marc Brillantes, Lucy Glover, Zdeněk Paris, Brian T. Chait, Mark C. Field, Michael P. Rout

## Abstract

In opistokhonts (animals and fungi), mRNA export to the cytoplasm is mediated by the Mex67/Mtr2 (NXF1/NXT1) heterodimer via the nuclear pore complex (NPC). In contrast to most nucleocytoplasmic transport, mRNA export requires ATP-dependent remodeling machinery, and in animals and fungi is Ran-independent. While most eukaryotes possess one Mex67 gene, trypanosomes have three distinct Mex67 paralogs, while retaining a single Mtr2 gene. We show here that these paralogs, TbMex67, TbMex67b and TbMex67L, have differing and non-redundant roles in RNA export. Specifically, TbMex67 and TbMex67b retain a canonical role in mRNA export, albeit associating with specific mRNA cohorts, but in contrast, TbMex67L is primarily involved in ribosome biogenesis. Together with the association of all Mex67 paralogs with the Ran machinery, these findings indicate significant departures in RNA export mechanisms in these divergent organisms, with implications for evolutionary origins and diversity in control of gene expression.

## Introduction

Controlling gene expression is an essential, complex, and highly regulated process, with many core mechanisms shared across the tree of life, reflecting their ancient origins. For eukaryotes many layers of expression control are exercised through variation in chromatin states, activation or repression of promoters and enhancers and, at the level of the transcript itself, through splicing, regulatory 5’ and 3’ untranslated regions (UTRs), translational efficiency and turnover (Daniel et al., 2014; Kaikkonen et al., 2011; Shaul, 2017; Wu, 1997). After transcription, processing and maturation, mRNA molecules must be translocated to the cytoplasm via nuclear pore complexes (NPCs) (Wozniak and Lusk, 2003) prior to translation (Okamura et al., 2015; Strambio-De-Castillia et al., 2010).

Transport through the NPC is mediated by factors that interact with a particular class of nucleoporins, namely phenylalanine-glycine (FG) dipeptide-rich proteins (termed FG-Nups). Directionality of most nucleocytoplasmic transport is provided by a gradient of predominantly RanGTP in the nucleus and RanGDP in the cytoplasm (Wente and Rout, 2010). However, bulk mRNA export in animals, fungi and plants is an ATP-dependent and Ran-independent process. In particular, an ATP-dependent DEAD box RNA helicase called Dbp5 (DDX19 in animals), and Gle1, an RNA export mediator, associate with the cytoplasmic-sided FG-repeat-containing Nup159 (Nup214 in animals) to remodel ribonucleoproteins (mRNPs) exiting the NPC (Adams et al., 2014; Alcazar-Roman et al., 2006; Hodge et al., 1999; Hodge et al., 2011; Montpetit et al., 2011; Noble et al., 2011; Weirich et al., 2004; Weirich et al., 2006). This step is required prior to dissociation of the heterodimeric RNA export Mex67:Mtr2 (NXF1:NXT1 in animals), which is then recycled into the nucleus, providing at least in part directionality and energy driving mRNA export (Katahira et al., 1999). Both Mex67 and Mtr2 have domains that interact specifically with FG-Nups: an NTF2-homologous domain formed by dimerization of Mtr2 with an Ntf2-like domain in Mex67; and a C-terminal UBA domain (Bayliss et al., 2002; Fribourg and Conti, 2003; Kang and Cullen, 1999; Katahira et al., 1999; Segref et al., 1997).

Most understandings of nucleocytoplasmic transport, NPC organization and roles in gene regulation stem from work in yeast and vertebrates, which are both members of the Opisthokonta, one of five major supergroups of the eukaryotic lineage (Adl et al., 2012; Baldauf, 2003). Less is known for other lineages despite significant reasons to suspect mechanistic divergence in RNA metabolism. Of these, trypanosomes represent both an extremely distant relative of animals/fungi and significant work has demonstrated very unusual transcriptional mechanisms. Trypanosomes lack canonical RNA polymerase II (Pol II) promoters for individual genes, instead relying on histone modifications and sequence specific promoters to drive polycistronic transcription of multiple genes with resolution to single mRNAs by *trans*-splicing and polyadenylation (Berriman et al., 2005; Cordon-Obras et al., 2022; Siegel et al., 2009). While mRNA stability is a key component of gene regulation in trypanosomes (Clayton, 2013), the role of mRNA export in regulating mRNA transcripts remains poorly characterized. Trypanosomes seem to bypass the surveillance system that prevents defective or unspliced mRNA from exiting the nucleus as blocking *trans*-splicing does not prevent defective or unspliced mRNA from exiting the nucleus (Goos et al., 2019). Nonetheless, although incompletely processed mRNA transcripts do exit the nucleus, they aggregate in granules in the cytoplasmic vicinity of the *Trypanosoma brucei* NPC (TbNPC) and are tethered there by an unknown mechanism (Goos et al., 2019). Moreover, as previously reported, the composition and architecture of the TbNPC indicates non-canonical mRNA export (Obado et al., 2016a).

Export factors Mex67 and Mtr2 are conserved in trypanosomes, and are termed TbMex67 and TbMtr2 respectively (Dostalova et al., 2013; Kramer et al., 2010; Schwede et al., 2009). TbMex67 (Tb927.11.2370) interacts directly with the NPC, TbMtr2, Ran, Ran-binding protein 1 (RBP1) and a GTPase activating protein (GAP) (Obado et al., 2016a), while recent evidence indicates that silencing TbMex67 or TbMtr2 compromises polyA RNA export (Dostalova et al., 2013) suggesting that TbMex67:TbMtr2 are *bona fide* mRNA export factors. TbMex67 and TbMtr2 also have roles in tRNA export (Hegedusova et al., 2019) and export of pre-ribosomal subunits (Buhlmann et al., 2015; Rink et al., 2019), analogous to opisthokonts (Chatterjee et al., 2017; Faza et al., 2012; Yao et al., 2008; Yao et al., 2007).

Here we report that trypanosomatids have diversified mRNA export to include additional divergent paralogs of Mex67. Only multicellular organisms are known to have additional Mex67 isoforms, some of which are expressed at lower levels, have tissue specificity, are developmentally regulated, and in most cases are generated as splice variants (Herold et al., 2000; Izaurralde, 2002; Jun et al., 2001; Kerkow et al., 2012; Yang et al., 2001). We show that the divergent trypanosome Mex67 paralogs also facilitate rRNA biogenesis and export, indicating a division of labor between these gene products. Coupled with a heavy reliance on post-transcriptional regulation, we suggest that mRNA export is an important step in regulating life cycle-specific mRNA transcripts in trypanosomes and that the RNA export factor Mex67 has RNA processing and assembly functions that can become separated from a role solely as a transport factor.

## Results

### Trypanosomatids possess three Mex67 paralogs

Divergence within the trypanosome NPC structure suggests the presence of distinct mRNA export mechanisms (Obado et al., 2016a). Specifically the NPC lacks the canonical mRNA Nup159/Gle1/Dbp5 remodeling platform (Nup214/Gle1/DDX19 in humans) (Adams et al., 2014; Fernandez-Martinez et al., 2016; Folkmann et al., 2011). To examine this pathway in greater detail we *in situ* GFP-tagged TbMtr2 and performed cryo-milling affinity capture to identify interacting proteins. Under permissive conditions TbMtr2 interacted with the entire NPC, similar to TbMex67 (Obado et al., 2016a). However, using more stringent conditions, a subset of trypanosome NPC nucleoporins (TbNups) (**Fig 1A**) were found, including the TbNup76 complex (TbNup76, 140, 149) and components of the outer ring complex (TbNup89, 132, 158). The TbNup76 complex is most likely analogous to the ScNup82 mRNA export platform. A connection between the TbNup76 complex and the trypanosome NPC outer ring is also probably conserved with opisthokonts, where the mRNA export platform (ScNup82/HsNup88 complexes) is also tethered to the NPC by the outer ring complex (Fernandez-Martinez et al., 2016). In addition, TbMtr2 co-purified with the entire Ran cycle apparatus, specifically the Ran GTPase, Ran-binding protein 1 (RanBP1) and a TBC-domain containing GTPase-activating protein (TbGAP) (Tb927.10.7680) (**Fig 1A**) (Gabernet-Castello et al., 2013; Goos et al., 2017; Obado et al., 2016a). Tb927.10.7680 was designated as TBC_Root_A by us earlier because of divergence from other Rab GAPs and specific presence within the TBC phylogeny root (Gabernet-Castello et al., 2013), consistent with a lineage-specific mRNA export pathway as well as the presence of a novel regulator (Obado et al., 2016a). Moreover, the close relationship between Ran and the Rab GTPases may indicate that originally these proteins utilized a GAP paralog, which has been replaced or diverged in higher eukaryotes so that the TBC domain is no longer detectable.

**Figure 1.**
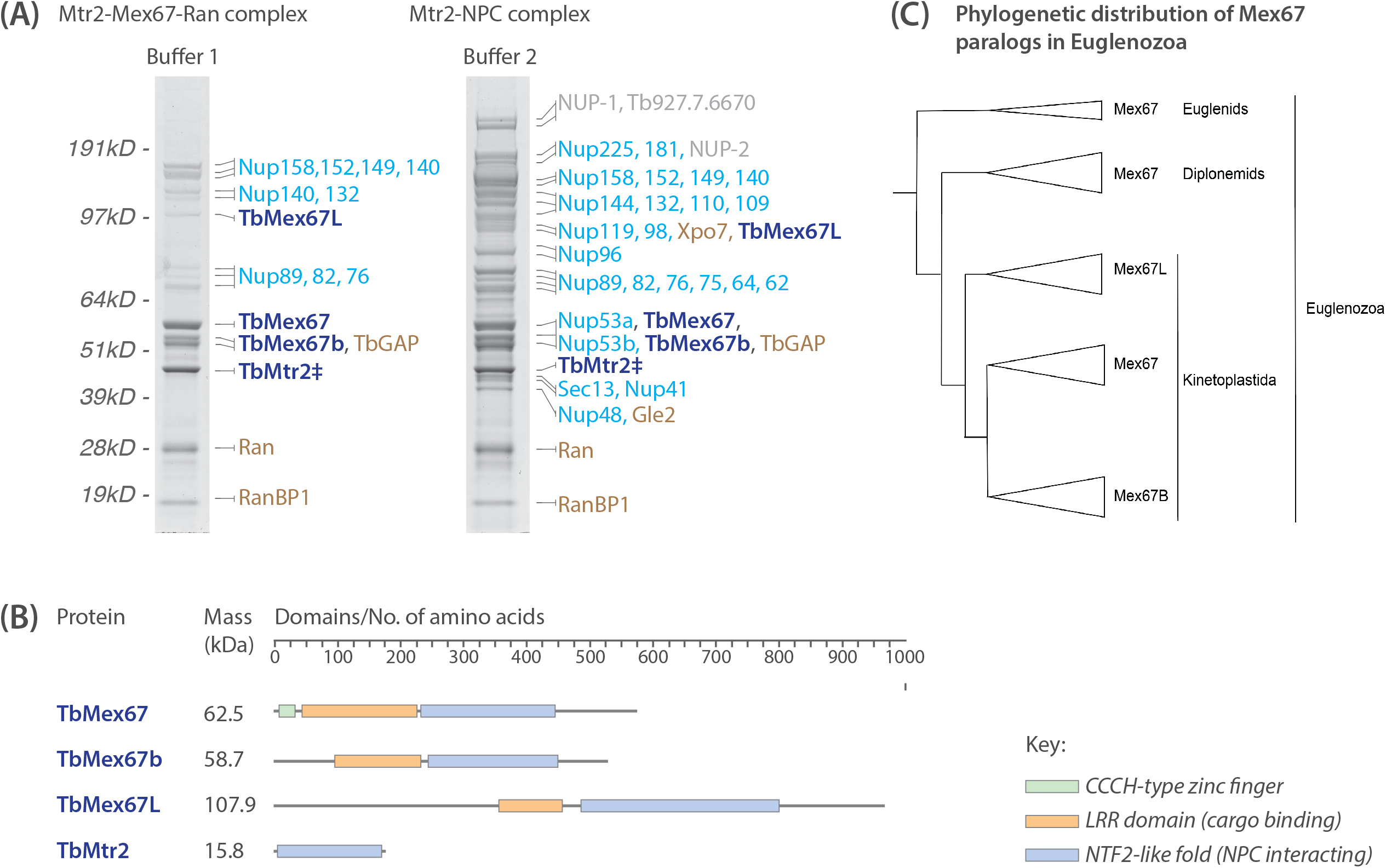
Affinity capture of the TbMtr2 and identification of new Mex67 paralogs. (A) Using the GFP tagged TbMtr2 (marked with a ‡), under high stringency biochemical conditions (Buffer 1), we affinity isolated structural components of the NPC (sky blue), transport factors connected to the GTPase Ran cycle (orange), and TbMex67 (navy blue) with which TbMtr2 is known to form a heterodimer (Dostalova et al., 2013). In addition we discovered interaction with two putative TbMex67 paralogs which we termed TbMex67b and TbMex67L (for like or large) denoted in navy blue like TbMex67. Under low stringency conditions (Buffer 2), we affinity isolated the entire trypanosome NPC as well as exportin 7 (Xpo7), the RNA export component Gle2 (Blevins et al., 2003; Murphy et al., 1996; Pritchard et al., 1999) and the lamin proteins NUP-1 and NUP-2 (DuBois et al., 2012; Maishman et al., 2016), and a hypothetical protein (gray). Affinity isolates were resolved by SDS-PAGE and visualized by Coomassie staining. Proteins bands were excised and identified by mass spectrometry. (B) A schematic of the domains that are present on each protein and highlights the difference between them. (C) A phylogenetic tree showing the distribution of the TbMex67 paralogs within Euglenozoa.

Significantly, TbMtr2 interacted with TbMex67 as expected, but also two additional proteins, Tb927.11.2340 (58kDa) and Tb927.10.2060 (108kDa). We designate these TbMex67b and TbMex67L (for TbMex67-Like) respectively, as both interact with TbMtr2 and contain characteristic LRR and NTF2-like domains, defining architectural features of Mex67 (Herold et al., 2000; Segref et al., 1997) (**Fig. 1B**) as predicted by AlphaFold (Jumper et al., 2021).

### Trypanosome Mex67 paralogs are distinct

TbMex67 has a CCCH-type zinc finger that differentiates it from opisthokont Mex67 orthologs, but which is essential for function as an mRNA export factor (Dostalova et al., 2013; Schwede et al., 2009). Additionally, TbMex67 has an LRR and a NTF2-like domain (Dostalova et al., 2013; Rink and Williams, 2019) (**Fig. 1B**). TbMex67b lacks the CCCH zinc finger, but retains the LRR and NTF2-like domains (**Fig. 1B**). TbMex67L is more divergent and is a significantly larger protein than TbMex67 and TbMex67b, due to an extended N-terminal domain consisting of three predicted alpha-helices on the AlphaFold database (Jumper et al., 2021). Nonetheless, TbMex67L also contains the characteristic LRR and NTF2-like domains, typical of opisthokont Mex67 proteins (**Fig. 1B**).

The genes encoding TbMex67 and TbMex67b are only 5kb apart on chromosome 11, indicative of a gene duplication event. Moreover, the chromosomal environment, including the synteny of these genes, is conserved throughout the kinetoplastids. Phylogenetic reconstruction across all kinetoplastids indicates that TbMex67 and TbMex67b are much more closely related than they are to TbMex67L, as expected (**Fig 1C**). Significantly, orthologs of TbMex67 and TbMex67b are recovered from all kinetoplastid genomes, including the free-living bodonid, *Bodo saltans*, implying that duplication predates the origin of parasitic trypanosomatids. By contrast, Mex67L is absent in the *B. saltans* genome, albeit retaining a presence within all other kinetoplastids, suggesting that Mex67L has a more recent origin compared with TbMex67 and TbMex67b.

### TbMex67 paralogs are differentially localized

The significant divergence in sequence and structure of the three Mex67 homologs led us to hypothesize that they have distinct functions. We *in situ* GFP-tagged the three TbMex67 paralogs in both procyclic forms (PCFs - insect stage parasites) and bloodstream forms (BSFs - vertebrate stage parasites) to determine their localizations by fluorescence microscopy (**Fig. 2A**). Similar to TbMtr2 localization in PCFs, TbMex67 and TbMex67b strongly localize to the nuclear envelope in puncta reminiscent of nucleoporins and other NPC-associated proteins (DeGrasse et al., 2009), in both BSFs and PCFs (**Fig. 2A**). They also localize around the nucleolus (**Fig. 2A**), as previously observed for TbMex67 in PCFs (Kramer et al., 2010). Yeast Mex67 is similarly concentrated at NPCs as well as having a diffuse localization in the nucleoplasm (Oeffinger et al., 2007b). In sharp contrast, TbMex67L is primarily nucleolar with no discernable peripheral nuclear punctate staining, indicating a significant difference in its functional role (**Fig. 2A)**.

**Figure 2.**
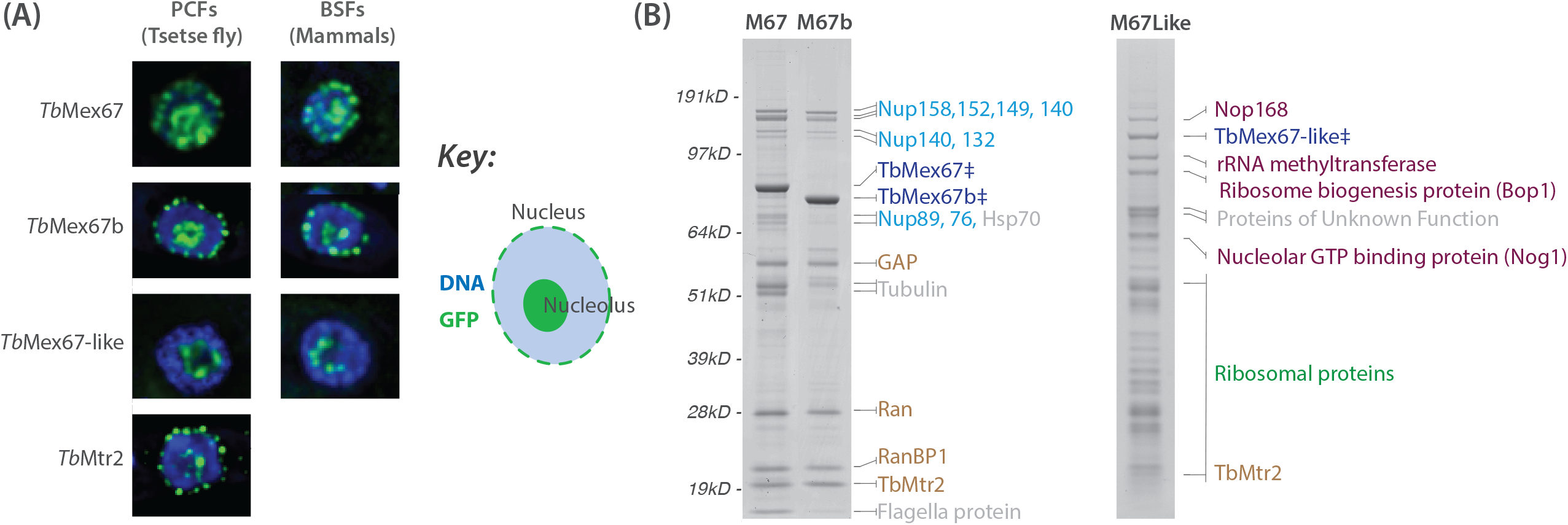
GFP tagging of the TbMex67 paralogs and their protein interactomes. (A) We in situ GFP tagged TbMex67, TbMex67b and TbMex67L in both the procyclic forms (PCFs) that infect tsetse flies and the bloodstream forms (BSFs) that infect vertebrates. The TbMtr2-GFP from PCFs is shown for comparison. We found that except for TbMex67L, other paralogs and TbMtr2 localize to nuclear pore complexes (NPCs) at the nuclear envelope as well as the nucleoplasm albeit predominantly around the nucleolus. (B) We performed affinity capture using each GFP tagged paralog in PCFs (marked with a ‡), and discovered that TbMex67 and TbMex67b interact predominantly with the NPC, whilst TbMex67L interacts only with ribosome biogenesis proteins, possibly a reflection of its nucleolar only localization.

### TbMex67 paralogs have distinct protein interactomes

We next investigated the potential differential roles of the three Mex67s by determining the interactomes for each in PCFs. Affinity captures of TbMex67 and TbMex67b co-purified similar protein cohorts. (**Fig 2B**). Both associate with Mtr2 as expected, given that they both co-purify with affinity captures of Mtr2 (**Fig 1A and 2B**). Both paralogs also associate with a specific subset of nucleoporins. Among these are FG-Nups TbNup140 and TbNup149 and the associated β-propeller, coiled-coil TbNup76, which together comprise the TbNup76 complex attached to the NPC outer rings towards the periphery on both the nuclear and cytoplasmic sides of the NPC; accordingly, we also found outer ring proteins TbNup152, TbNup132, TbNup89 and TbNup158, (DeGrasse et al., 2009; Obado et al., 2016a). We suggest that TbMex67 and TbMex67b interact at the NPC with the three FG-Nups (140, 149 and 158), and indirectly with Nup76 via its attachment to the outer ring. This robust association with nucleoporins is consistent with the localization of both proteins to the NPC at the nuclear periphery (**Fig 2A**). TbMex67 and TbMex67b also both interact with the Ran transport factor system: Ran, RanBP1 and the putative RanGAP, TBC-RootA, as seen for Mtr2 (Gabernet-Castello et al., 2013; Obado et al., 2016a). Collectively, these data indicate that both Mex67 paralogs function as nuclear transporters; as TbMex67 functions as an mRNA exporter (Dostalova et al., 2013), we suggest that this function is shared by both proteins.

The TbMex67L interactome contains Mtr2 as expected (**Fig 1A and 2B**), consistent with a constitutive function for the Mex67L:Mtr2. However, there is no concordance between the TbMex67L and TbMex67/TbMex67b interactomes, but rather a subset of ribosomal proteins and nucleolar-associated proteins involved in ribosomal biogenesis were recovered with TbMex67L. Amongst these, Nop168 is nucleolar localized (Dean et al., 2017) and trypanosome specific, while Nog1 is a widely conserved nucleolar GTPase (Dean et al., 2017), co-ordinating ribosome formation (Klingauf-Nerurkar et al., 2020). Nop89, also nucleolar localized (Dean et al., 2017), is required for synthesis of the 28S and 5.8S ribosomal RNAs (rRNAs) and thus large ribosome subunits (Pestov et al., 2001). Significantly, no nucleoporins or Ran transport factor system proteins were identified, consistent with an absence from the nuclear periphery. Instead, localization to the nucleolus and an association with ribosomal proteins and nucleolar localized ribosomal biogenesis proteins strongly indicates a role in ribosomal biogenesis (Faza et al., 2012; Fribourg et al., 2001; Herold et al., 2000; Katahira et al., 1999; Kent et al., 1999; Sarkar et al., 2016; Yao et al., 2008; Yao et al., 2007).

### All TbMex67 paralogs are required for viability

We used RNA interference (RNAi) to test for essentiality and functional redundancy between the three TbMex67 paralogs. Silencing each paralog results in severe loss of fitness in BSFs, suggesting independent essentiality (**Fig. 3**). TbMex67 is essential in PCFs (Buhlmann et al., 2015; Dostalova et al., 2013; Hegedusova et al., 2019), and our RNAi result recapitulates this phenotype in PCFs using the same cell line as a control (Hegedusova et al., 2019) (**Fig. 3**), but silencing TbMex67b or TbMex67L resulted in a gradual loss of fitness over several replicative cycles in PCFs, suggestive of a cumulative and possible secondary impact (**Fig. 3**).

**Figure 3.**
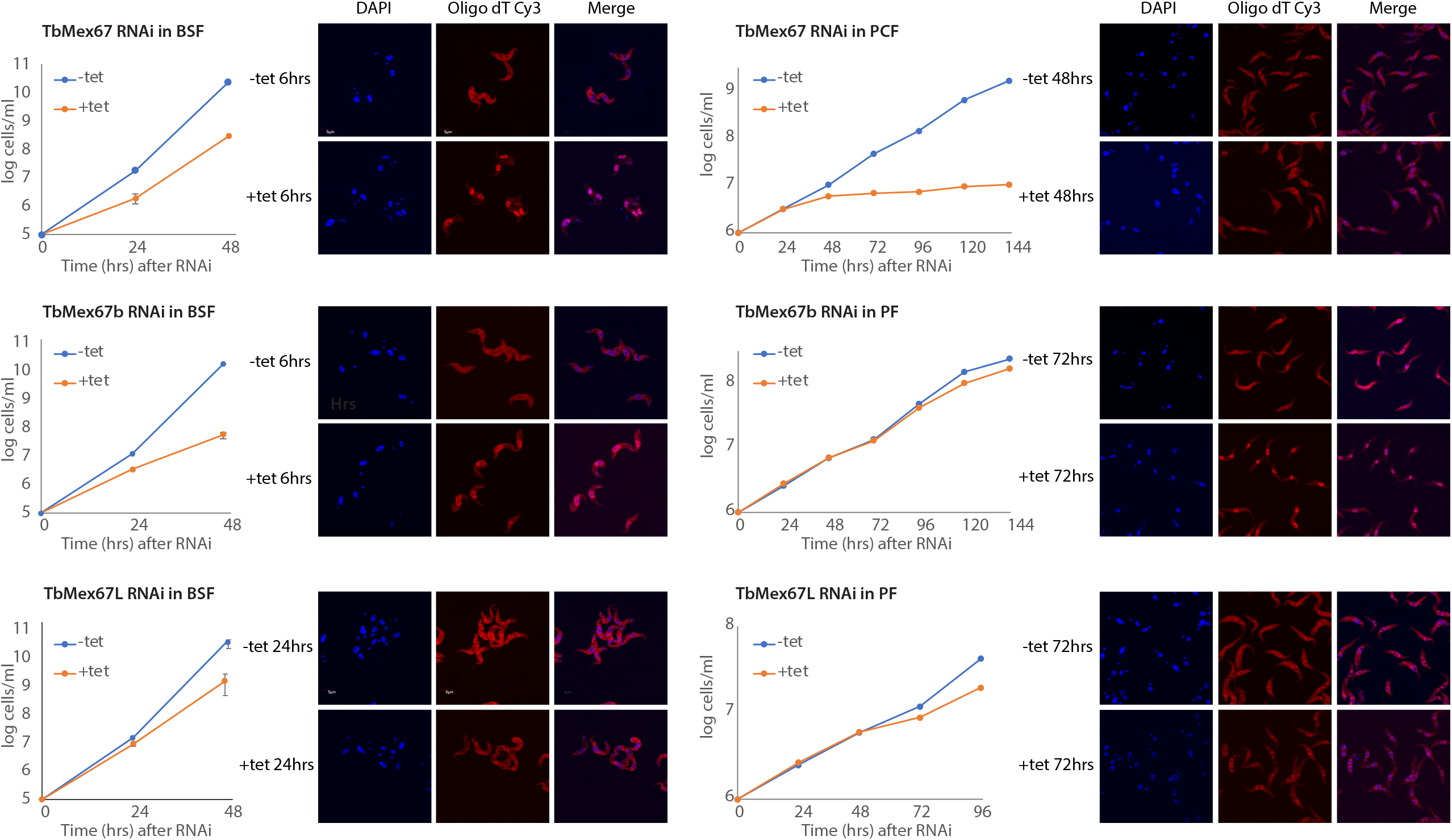
RNA interference knockdown of the Mex67 paralogs and poly A FISH to determine role in RNA export. To assess the role of the TbMex67 paralogs on mRNA export, we performed RNA FISH after induction of RNAi silencing for each individual paralog. The loss of fitness in BSFs occurs much faster and is accompanied by a rapid accumulation of mRNA in the nucleus of TbMex67 and TbMex67b cell lines, but not TbMex67L in which there is no mRNA accumulation indicating it is not involved in RNA export. In PCFs there was a rapid loss of fitness in TbMex67 cell lines post RNAi induction and accumulation of mRNA in the nucleus. Similarly, there was accumulation of mRNA in the nuclei of the TbMex67b cell lines but at a much slower rate. Likewise, RNAi loss of fitness was very gradual post RNAi induction in TbMex67L cell lines. However, unlike in BSFs, over time, there is a perturbation of mRNA export exhibited as a halo around the nuclei of the TbMex67 cell line. This is under investigation.

### Only TbMex67 and TbMex67b are required for mRNA export

In addition to testing for viability, we used polyA FISH (fluorescence *in situ* hybridization) to detect a function in the export of polyadenylated mRNA. Silencing TbMex67 or TbMex67b in BSFs leads to accumulation of polyA in the nucleus and depletion from the cytoplasm, with probable accumulation around the cytoplasmic periphery of the nucleus (**Fig. 3**). This, in conjunction with location and binding partners, indicates that TbMex67 and TbMex67b are *bona fide* mRNA export components (Buhlmann et al., 2015; Dostalova et al., 2013; Hegedusova et al., 2019).

Loss of fitness in PCFs following RNAi knockdown varies considerably between TbMex67 and TbMex67b. There is significant nuclear accumulation of polyA in TbMex67 silenced cells shortly after RNAi silencing (**Fig. 3**) (Dostalova et al., 2013), but this is much slower following RNAi knockdown of TbMex67b, suggestive of cumulative RNAi effects over time.

Further, silencing TbMex67L results in distinct mRNA localizations in the two life stages. There is no nuclear accumulation of mRNA in BSFs, correlating with having no role in nuclear export, but in contrast in PCFs, there is a gradual accumulation of mRNA on the periphery of the nuclear envelope (**Fig. 3**); this accumulation is reminiscent of the nuclear periphery granules observed to form upon inhibition of trans-splicing but is also a possible secondary effect due to loss of ribosomal assembly and hence protein synthesis (Goos et al., 2019; Kramer et al., 2012). Overall, it is clear that TbMex67L is primarily involved in non-mRNA related functions.

### Differential RNA associations of TbMex67 and TbMex67b

Additionally to identifying proteins forming stable interactions with the TbMex67 paralogs, we affinity isolated TbMex67 and TbMex67b from PCFs under conditions ensuring maximal preservation of RNA-protein interactions (Aitchison et al., 1996; Domanski et al., 2016; Oeffinger et al., 2007a; Oeffinger et al., 2007b; Oeffinger et al., 2009) (see methods). As expected TbMex67 and TbMex67b associate strongly with a large number of mRNA species, as well as rRNA (also expected as opisthokont Mex67 plays a role in ribosomal subunit export) (Faza et al., 2012; Sarkar et al., 2016; Yao et al., 2008; Yao et al., 2007). Isolation of mRNA, NPC proteins and the Ran GTPase system with TbMex67 and TbMex67b indicates that we are capturing at least some of the nucleoplasmic, export and cytoplasmic pools of the complexes formed by these paralogs (Okamura et al., 2015). We found that the majority of mRNA cargo isolated are mostly in common, including abundant mRNA species such as elongation factor α1 (TEF1), which associates with a specific subset of mRNAs in *T. cruzi* (Alves et al., 2015), and pyruvate phosphate dikinase (PPDK), a glycolytic enzyme specific to PCFs (Vertommen et al., 2008) (**Fig. 4**). However, we found a very significant enrichment of a subset of mRNA species unique to TbMex67 (**Fig. 4**). Moreover, the mRNA species specifically enriched in TbMex67 affinity isolates are those that encode for RNA-binding proteins (RBPs) with DRBD12 (Tb927.7.5380), RBP26 (Tb927.7.3730) and ZC3H46 (Tb927.11.16550) being most abundant mRNA species in this cohort (**Fig. 4**). RBP26 and DRBD12 are amongst six mRNA transcripts that have been observed to be upregulated after the induction of RPB6, the master regulator of the transformation of non-infectious PCFs into infectious metacyclic forms (Shi et al., 2018). ZC3H46 and DRBD12 have previously been demonstrated to negatively affect gene expression in a screen that selects for proteins that impair expression of a lethal phosphoglycerate kinase B reporter (Lueong et al., 2016). Also, inhibition of DRBD12 by RNAi followed by microarray analysis of the transcriptome leads to upregulation of mRNAs containing AU-rich elements (ARE) in their 3’ UTR and overexpression led to down-regulation of ARE-containing transcripts (Najafabadi et al., 2013). It has recently been shown that TbMex67 and TbMtr2 also associate with RBPs themselves to regulate the export of specific cohorts of mRNA. Downregulation of DRBD18 results in partial accumulation of TbMex67 and TbMtr2 and perturbation of the export a subset of mRNAs out of the nucleus (Mishra et al., 2021).

**Figure 4.**
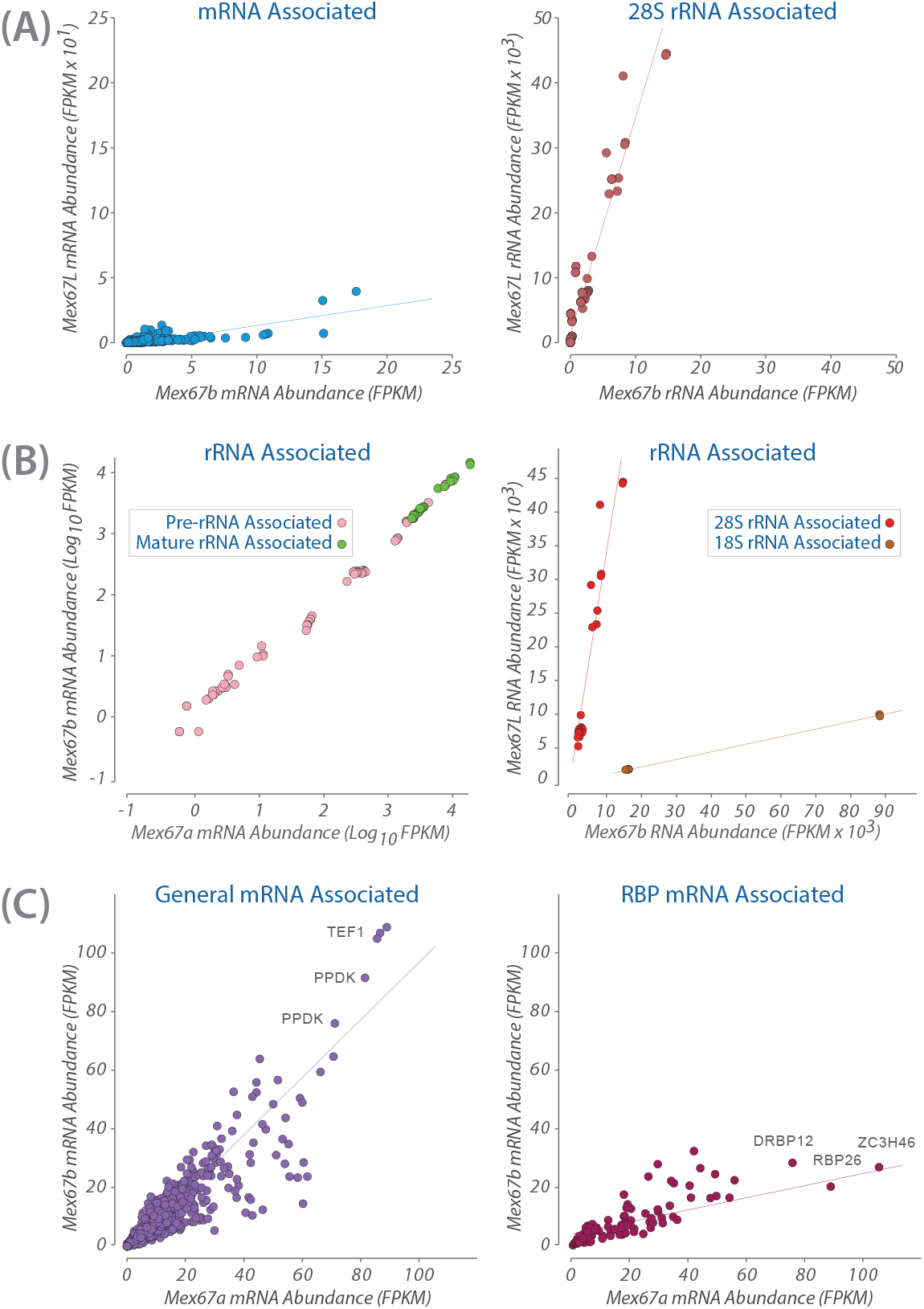
Affinity capture and identification of the RNA cargo of each TbMex67 paralogs. We affinity captured each TbMex67 paralog in PCFs and then extracted interacting RNA and performed Illumina sequencing to determine the which RNA species each paralog interacts with. (A) A comparison of RNA cargo between the RNA exporting TbMex67b versus the ribosome biogenesis TbMex67L. TbMex67b interacts with mRNA whereas TbMex67L does not. Instead TbMex67L interacts with 28s rRNA. (B) We compared rRNA sequences that associate with all three TbMex67 paralogs. We find that once again TbMex67L interacts with 28s rRNA, and when compared to TbMex67b which interacts predominantly with 18S rRNA. Both TbMex67 and TbMex67b interact equally with 18S rRNA. (C) TbMex67 and TbMex67b share common mRNA cargo. However, there is a significant enrichment of mRNA encoding RNA binding proteins (RBPs) on TbMex67 as compared with TbMex67b, suggesting a role in control of gene expression given the role of RBPs in the post transcriptional regulation of genes in trypanosomes (Clayton, 2013).

**Figure 5.**
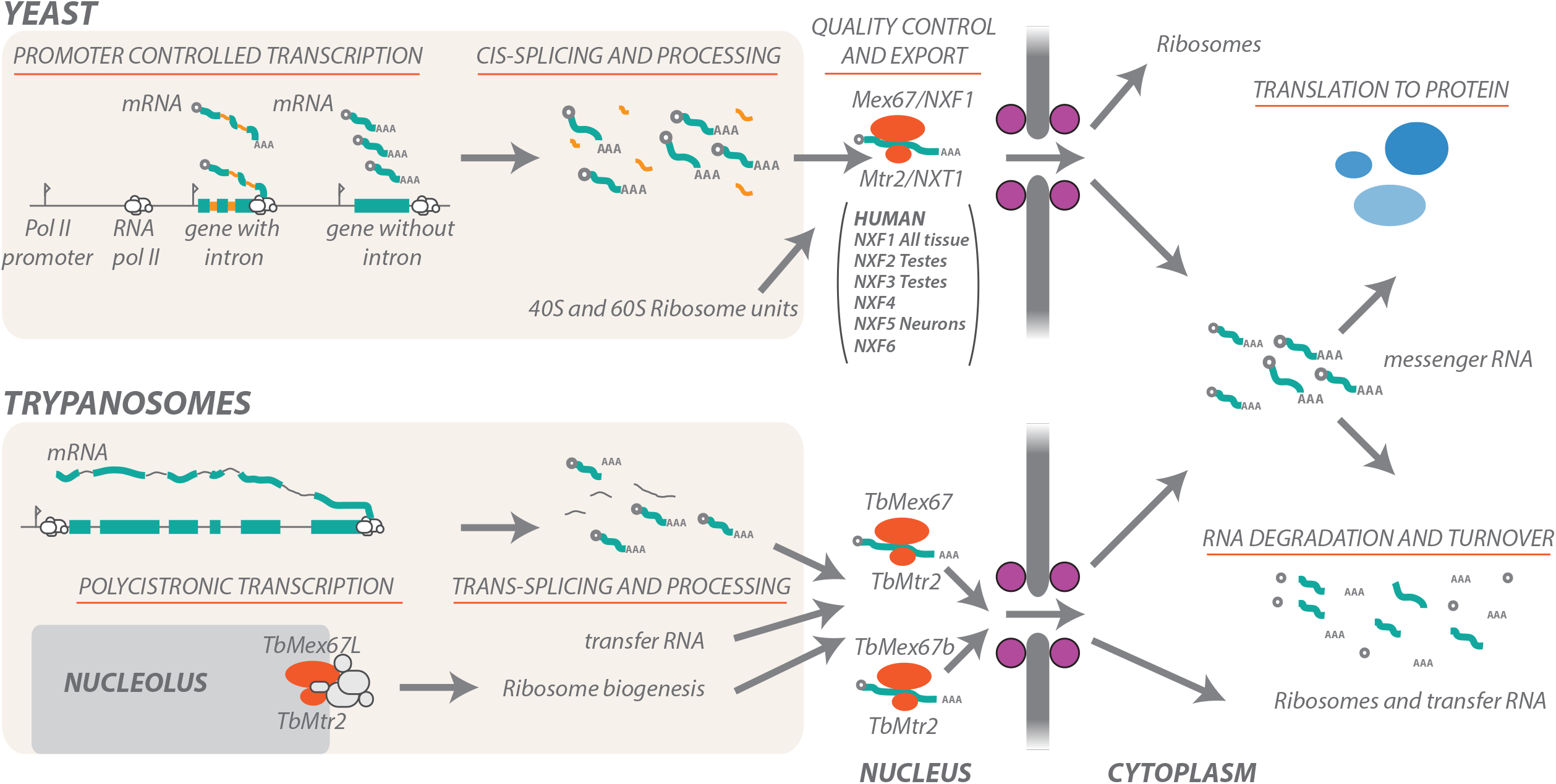
Model of the roles of the three TbMex67 paralogs and RNA export in comparison to opithokonts. In most eukaryotes, each gene is has a dedicated Polymerase II (Pol II) promoter. Genes are transcribed, capped, introns spliced out, polyadenylated and export occurs co-transcriptionally, with the basket playing a prominent role in quality control prior to mRNA export. Trypanosomes have histone marks and single sequence specific promoters for each polycistronic transcription unit as opposed to individual gene promoters (Berriman et al., 2005; Cordon-Obras et al., 2022; Siegel et al., 2009). Transcription is polycistronic with individual mRNA resolved by trans splicing of a splice leader sequence common to every single mRNA and then poly adenylated (comprehensively reviewed in (Kramer, 2021)). Thus, mRNA export has an important role in gene regulation in trypanosomes. mRNA is exported by the Mex67/Mtr2 heterodimer (Katahira et al., 1999). Multicellular organisms have additional tissue specific Mex67 paralogs known as NXFs (Herold et al., 2000). Trypanosomes are the only single celled organism thus far to be shown to have 3 distinct Mex67 paralogs, one of which is involved in rRNA biogenesis and not mRNA export. Additionallky, unlike opisthokonts, tRNA export is mediated by TbMex67/TbMtr2 (Hegedusova et al., 2019).

TbMex67 and TbMex67b associate equally with rRNA (**Fig. 4**) and are presumably involved in their biogenesis and export as has been demonstrated in trypanosomes (Buhlmann et al., 2015; Rink et al., 2019; Rink and Williams, 2019). Whilst it is uncertain whether Mex67/Mtr2 are involved in ribosome biogenesis in opisthokonts, they are known to export 60S and 40S ribosomal units (Faza et al., 2012; Yao et al., 2007).

### TbMex67L performs non-canonical functions

We also affinity captured TbMex67L and isolated the associated RNA in a similar manner to TbMex67 and TbMex67b. Following RNA-Seq, we found that TbMex67L interacts exclusively with rRNA and not mRNA, and specifically 60S, the large ribosomal subunit RNA (**Fig. 4**). Taken together with its protein interactome, we conclude that TbMex67L is limited to ribosome biogenesis and not nucleocytoplasmic export of RNA cargos, despite the RNAi phenotype in PCFs where there appears to be a mRNA halo around the nucleus (**Fig. 3**). A possibility for the RNAi phenotype may be a general result of collapsing protein synthesis and gradual loss of many RNA export functions. Hence trypanosomes possess an evolutionary diversification in Mex67 functions, to include a paralog with rRNA/ribosomal duties exclusively. Given that Mtr2 interacts with all three TbMex67 paralogs, its disruption likely leads to multiple pleiotropic effects that would impact the export of RNA from the nucleus as well as ribosome biogenesis.

## Discussion

Unusually for a unicellular organism trypanosomatids have diversified the ancestral Mex67 into three paralogs. All three share a common protein architecture and form heterodimers with the conserved Mex67 co-factor, Mtr2. Outside of the conserved core, each paralog has distinct features. TbMex67 uniquely has a well characterized N-terminal CCCH zinc finger motif essential for its function (Dostalova et al., 2013). All three possess LRR and NTF2 domains, characteristic of Mex67 in opisthokonts, and that act synergistically with the NTF2 domain to mediate interactions with FG-Nups (Fribourg et al., 2001; Fribourg and Conti, 2003). TbMex67L has an N-terminal extension accounting for its significantly larger molecular weight, which is predicted to be composed primarily of three bundled α-helices.

The Mex67 paralogs also have distinct functions. TbMex67 and TbMex67b are bona fide mRNA exporters since they interact with the NPC, associate with mRNA and rRNA and their RNAi knockdown results in accumulation of mRNA in the nucleus. Additionally, Mex67b was recently identified in a proteomic screen in *T. cruzi* using TcSub2 as the affinity handle, strongly supporting its role as being involved in mRNA export (Inoue et al., 2022). Although our interactome doesn’t have an abundance of RNA binding proteins, we detect poly A binding protein at sub stoichiometric levels, highlighting the dynamism of the system. Additionally, it has been shown in opisthokonts that Mex67 behaves as a mobile nucleoporin the majority of which is not bound to mRNA in the nucleus (Derrer et al., 2019). Thus what we observe in our interactome may be the steady state, with messenger ribonucleoparticles interacting weakly and in a fast transient manner. It is well documented in opisthokonts that Mex67/NXF1 are involved in the export of rRNA through the Crm1/XPO1 – NMD3 pathway (reviewed in (Kohler and Hurt, 2007)). Similarly, TbMex67 participates in the NMD3 pathway in trypanosomes, suggesting a level of evolutionary conservation of this role (Buhlmann et al., 2015). We suggest that TbMex67 and TbMex67b also have a role in 40S subunit export, as evidenced by their interaction with 18S rRNA. TbMex67 may also function in 60S export in trypanosomes via interaction with the 5S RNP (Rink and Williams, 2019). Nevertheless, there are important differences in the mRNAs that TbMex67 and TbMex67b bind, most notably that TbMex67 has a significantly higher association with mRNAs encoding RBPs (**Fig. 4**), suggestive of a role in gene regulation in PCFs.

TbMex67L has no obvious role in mRNA or ribosomal subunit export, as it lacks interactions with NPC-associated proteins. Instead, several lines of evidence indicate that TbMex67L operates in rRNA maturation and ribosomal assembly. TbMex67L interacts with several proteins involved in ribosome biogenesis such as Bop1 and Nog1. This functional distinction is curious, as TbMex67L retains both the Mtr2-interacting domain and an interaction with Mtr2, which together form an obligate FG-Nup repeat interacting region in other Mex67 homologs. We speculate that weak interactions with FG repeats may have been retained to allow a lower degree of FG-Nup association than we can currently detect, sufficient to aid in import of the heterodimer into the nucleus so that it can access the nucleolus. This functional distinction is, to our knowledge, unique amongst Mex67 homologs (Herold et al., 2000), but as Mex67 usually has two important roles, RNA processing/assembly to generate a mature RNP (Rink et al., 2019) and then export of the mature RNP out of the nucleus.

Multicellular organisms have closely related isoforms of Mex67 or NXF, some of which are generated as splice variants (Herold et al., 2000; Izaurralde, 2002; Jun et al., 2001; Kerkow et al., 2012; Yang et al., 2001). Humans have six NXF gene products, namely NXF1 to NXF6 (Herold et al., 2000). NXF1 (Mex67) is found in all tissues and thus considered the global mRNA exporter in metazoa whilst NXF2, NXF3, NXF5 and NXF7 have tissue specific functions (Herold et al., 2000). NXF2 is expressed in testes and neurons (Lai et al., 2006; Zhang et al., 2007), whilst NXF3 is expressed mainly in testes (Yang et al., 2001). NXF5 localizes to neurons and associates with translating ribosomes, stress granules and P-bodies(Alber et al., 2007; Jun et al., 2001; Katahira et al., 2008; Lai et al., 2006; Vanmarsenille et al., 2013). NXF1 and NXF2 predominantly localize to the nucleoplasm and display mRNA export activities, whilst NXF3 and NXF5 are mainly cytoplasmic, highlighting their functional differences (Herold et al., 2000; Tan et al., 2005). Importantly, NXF1, 2 and 3 form a heterodimer with NXT1 (Mtr2) which facilitates NPC localization and translocation (Izaurralde, 2002; Katahira et al., 1999; Santos-Rosa et al., 1998; Suyama et al., 2000; Wiegand et al., 2002). In humans, NXT1 has a paralog termed NXT2, which is generated as two splice variants, and can form a heterodimer with NXF1 *in vitro* (Herold et al., 2000).

Eukaryotic gene expression shares many ancestral properties with prokaryotic ancestors, albeit with additionally species-specific proteins and novel pathways arising to address the specific evolutionary adaptations of each organism. Trypanosomes have sculpted their mRNA processing/export pathways, probably as a consequence of polycistronic transcription and possibly also as a response to PolI transcription of protein encoding mRNAs. In trypanosome mRNA export and processing are Ran-dependent, a fundamental shift in how the pathway is controlled when compared with the canonical pathway in animals, fungi, plants and most other lineages. Trypanosomes have generally provided evolutionary lessons in the plasticity of conserved eukaryotic machineries, given other examples of highly distinct seemingly kinetoplastid functions such as an atypical lamina, nuclear basket and divergent kinetochores.

## Methods

### Cell Culture

*T. brucei* procyclic Lister 427 strain cells were cultured in SDM-79, supplemented with 10% fetal bovine serum as previously described (DeGrasse et al., 2009). Expression of plasmid constructs was maintained using Hygromycin B at 30 µg/ml.

### In Situ Genomic Tagging

All proteins tagged in this study used the pMOTag4G tagging vectors (Oberholzer et al., 2006) as previously described (DeGrasse et al., 2009; Obado et al., 2016a).

### Fluorescence Microscopy and RNA FISH

GFP-tagged cell lines were harvested and fixed for 10 mins in a final concentration of 2% paraformaldehyde. Fixed cells were then washed in 1xPBS and visualized as previously described (Obado et al., 2016a). RNA FISH was performed as previously described (Hegedusova et al., 2019). Briefly, 1 × 10^7^ cells were harvested and washed with PBS. Cells were resuspended in 4% paraformaldehyde/PBS solution and fixed to poly-l-lysine coated microscope slides for 30 min and dehydrated using an increasing gradient of ethanol concentrations (50, 80 and 100%, for 3 min each). Permeabilized cells were pre-hybridized with hybridization solution (2% BSA, 5× Denhardt’s solution, 4× SSC, 5% dextran sulphate, 35% deionized formamide, 10 U/ml RNase inhibitor), for 2 hrs, then incubated overnight at room temperature in a humid chamber in the presence of 10 ng/μl Cy3-labeled oligonucleotide probes in hybridization solution. Slides were then washed for 10 min, once with 4× SSC with 35% deionized formamide, then one wash each with 2× SSC and 1× SSC. Finally, the slides were mounted with mounting medium supplemented with 4′,6-diamino-2-phenylindole dihydrochloride (DAPI). Images were taken with confocal microscope Olympus FluoView™ FV1000 and analyzed using Fluoview and ImageJ (NIH) software. FISH data were quantified as described previously (Hegedusova et al., 2019).

### Affinity Isolation and purification of TbMex67 paralog protein interactome

Affinity capture of the TbMex67 paralogs was performed as previously described (Obado et al., 2016a). Briefly, trypanosomes were grown to a density of between 2.5 × 10^7^ cells per ml. Parasites were harvested by centrifugation, washed in 1xPBS with protease inhibitors and 10mM dithiothreitol, and flash frozen in liquid nitrogen to preserve protein:protein interactions as close as they were at time of freezing as possible. Cells were cryomilled into a fine grindate in a planetary ball mill (Retsch). For a very detailed protocol, refer to our methods paper (Obado et al., 2016b), or the National Center for Dynamic Interactome Research website (www.NCDIR.org/protocols). Cryomilled cellular materials were resuspended extraction buffers containing a protease inhibitor cocktail without EDTA (Roche), and clarified by centrifugation (20,000 x *g*) for 10 min at 4°C (Obado et al., 2016a; Obado et al., 2016b). Clarified lysates were incubated with magnetic beads conjugated with polyclonal anti-GFP llama antibodies on a rotator for 1 hr at 4°C. The magnetic beads were harvested by magnetization (Dynal) and washed three times with extraction buffer prior to elution with 2% SDS/40 mM Tris pH 8.0 for protein.

### Identification of protein interactome by Mass Spectrometry

Briefly, protein bands were excised from acrylamide gels and destained using 50% acetonitrile, 40% water, and 10% ammonium bicarbonate (v/v/w). Gel pieces were dried and resuspended in trypsin digestion buffer; 50 mM ammonium bicarbonate, pH 7.5, 10% acetonitrile, and 0.1–2 ug sequence-grade trypsin, depending on protein band intensity. Digestion was carried out at 37°C for 6 h prior to peptide extraction using C18 beads (POROS) in 2% TFA (trifluoroacetic acid) and 5% formamide. Extracted peptides were washed in 0.1% acetic acid (ESI) and analyzed on a LTQ Velos (ESI) (Thermo).

### Affinity Isolation and purification of TbMex67 paralog RNA cargo

Affinity capture was performed as described above, with the following modifications. We maximized RNA yield using ultrafast affinity capture with using our very high affinity double nanobody with a K_d_ of 36pM (Fridy et al., 2014). This has allowed us to complete affinity isolation experiments in ten minutes, minimizing degradation, and spurious non-specific interaction issues that plague lengthy incubation times. To minimize background, we incubated our magnetic beads used for affinity capture with poly dI-dC (deoxy-inosinic deoxy-cytidilic acid (Sigma) as a proven blocking agent to reduce non-specific RNA binding (Umaer and Williams, 2015), especially of abundant RNA species such as rRNAs. Rather than elute magnetic beads with SDS, RNA was extracted using the Qiagen RNA extraction kit. The magnetic beads used for affinity capture were incubated with Buffer RLT for 10 minutes and then RNA extracted using manufacturers protocols (Giagen). Buffer for protein interactome of TbMex67 and TbMex67b was 20mM HEPES, 250mM NaCl, 10uM MgCl_2_ and 0.5% Triton; 20mM HEPES, 150mM NaCl, 10uM MgCl_2_ and 0.5% Triton for RNA extraction. With TbMex67L 20mM HEPES, 20mM NaCl, 50mM tri-sodium citrate and 0.5% Triton for both protein and RNA extraction. We observed higher stoichiometries of RNA binding proteins, especially poly A binding protein 2 (PAB2), which were previously in sub stoichiometric amounts in our LC-MS analyses under the above biochemical conditions. We also tested stabilization of the TbMex67 complexes using glutaraldehyde, which was added to our affinity capture buffer, prior to resuspending our frozen cell grindate. We performed RNA-Seq using both glutaraldehyde stabilized and non-fixed samples (see methods) and saw no noticeable difference in RNA cargo. We used total cellular RNA as a control for read numbers and coverage versus enrichment of our affinity isolated pool, by normalizing the level of coverage of an observed RNA after purification to its total abundance in the lysate. This enabled us to determine comprehensive (transcriptome-wide) RNA associations with each life cycle stage through RNA-Seq.

We utilized Agilent RNA Pico chips to ensure the integrity and enrichment of RNA under various buffer conditions on small-scale purifications. Extracted RNA was then sequenced using The Rockefeller University Genomic Center”s Illumina HiSeq 2000.

### RNA interference and loss of function tests

RNAi was performed using the Horn lab”s 2T1 and PT1 cell lines (Alsford et al., 2011). Briefly, 1kb segments from each TbMex67 paralog was PCR amplified from *T. brucei* genomic DNA and cloned into the pRPa^iSL^ plasmid and transfected into 2T1 BSFs and PT1 PCFs. Cells were grown to a density of 2 × 10^7^ and induced using 1μg ml^-1^ of tetracycline. RNA FISH was performed at at the time points highlighted (**Fig. 3**).

### Secondary Structure Prediction and Structural Modeling

The three TbMex67 paralogs were analyzed for several secondary structure elements, Phyre2 (Kelley et al., 2015) and HHPred (Soding et al., 2005). 3D structures were modeled using the program I-TASSER (Yang and Zhang, 2015) and later using AlphaFold (Jumper et al., 2021).

## Notes

### Competing Interest Statement

The authors have declared no competing interest.

